# Impaired ureagenesis due to arginine-insensitive N-acetylglutamate synthase

**DOI:** 10.1101/425959

**Authors:** Parthasarathy Sonaimuthu, Emilee Senkevitch, Nantaporn Haskins, Prech Uapinyoying, Markey McNutt, Hiroki Morizono, Mendel Tuchman, Ljubica Caldovic

**Affiliations:** Center for Genetic Medicine Research, Children’s National Medical Center, Washington, DC, USA.; National Institute of Neurological Disorders and Stroke, National Institutes of Health, Bethesda, MD, USA.; Children’s Medical Center, UT Southwestern Medical Center, Dü, Dallas, TX, USA.

**Keywords:** urea cycle, ureagenesis, arginine, N-acetylglutamate synthase, N-acetylglutamate, regulation

## Abstract

The urea cycle protects the central nervous system from ammonia toxicity by converting ammonia to non-toxic urea. N-acetylglutamate synthase (NAGS) is an enzyme that catalyzes the formation of N-acetylglutamate (NAG), an allosteric activator of carbamylphosphate synthetase 1 (CPS1), the rate limiting enzyme of the urea cycle. Enzymatic activity of mammalian NAGS doubles in the presence of L-arginine but the physiological significance of NAGS activation by L-arginine is unknown. Previously, we have described the creation of a NAGS knockout (*Nags*^−/−^) mouse, which develops hyperammonemia without N-carbamylglutamate and L-citrulline supplementation (NCG+Cit). In order to investigate the effect of L-arginine on ureagenesis *in vivo*, we used adeno associated virus (AAV) mediated gene transfer to deliver either wild-type or E354A mutant mouse NAGS (mNAGS), which is not activated by L-arginine, to *Nags*^−/−^ mice. The ability of the E354A mNAGS mutant protein to rescue *Nags*^−/−^ mice was determined by measuring their activity on the voluntary wheel following NCG+Cit withdrawal. The *Nags*^−/−^ mice that received E354A mNAGS remained apparently healthy and active but had elevated plasma ammonia concentration despite similar expression levels of the E354A mNAGS and control wild-type NAGS proteins. The corresponding mutation in human NAGS (NP 694551.1:p.E360D) that abolishes binding and activation by L-arginine was also identified in a patient with hyperammonemia due to *NAGS* deficiency. Taken together, our results suggest that L-arginine binding to the NAGS enzyme is essential for normal ureagenesis.

## Introduction

Urea cycle functions in the liver to protect the central nervous system from the toxic effects of ammonia, a nitrogenous waste product of the catabolism of dietary and cellular proteins. Six urea cycle enzymes and two mitochondrial transporters collectively convert ammonia into urea, which is excreted through the kidneys [1]. Defects in any of the urea cycle enzymes or transporters lead to hyperammonemia, which can damage the brain and could be lethal if left untreated [2]. N-acetylglutamate synthase (NAGS; EC2.3.1.1) is a mitochondrial enzyme that catalyzes formation of N-acetylglutamate (NAG) from glutamate and acetyl coenzyme A [3-5]. NAG is an essential allosteric activator of carbamylphosphate synthetase 1 (CPS1; EC 6.3.4.16), the rate-limiting enzyme of the urea cycle [6, 7]. Enzymatic activity of mammalian NAGS doubles in the presence of L-arginine [3-5, 8, 9]; therefore, this amino acid could regulate the rate of urea production through modulation of NAGS activity to supply variable amounts of NAG for controlled activation of CPS1. This stipulation is supported by several lines of evidence. First, the rate of production of L-citrulline, a urea cycle intermediate, is proportional to the concentration of L-arginine in the portal blood, which correlates with dietary protein intake [10]. Second, intraperitoneal injections of L-arginine and L-glutamine lead to elevated NAG levels in rat liver [4]. Third, tracking of L-arginine metabolism in the liver mitochondria revealed that in addition to activating NAGS, L-arginine is converted to glutamate, which could activate NAGS through mass action as a substrate [11]. Finally, the NAG content in isolated liver mitochondria as well as hepatic content of NAG is higher in rats fed a high protein diet and this increase is paralleled by increased urea excretion [12-15]. The concentration of NAG in mouse and rat liver cells increases after a meal [16, 17] without need for new protein synthesis [18], suggesting that NAG is a short-term regulator of ureagenesis.

However, other evidence suggests that NAG, NAGS and L-arginine may not be regulators of urea synthesis. Although early measurements of the intramitochondrial NAG concentration indicated that it is comparable to the NAG activation constant for CPS1 [19], subsequent measurements indicated two to three times higher NAG levels, suggesting that most of the CPS1 molecules ought to be complexed with NAG and poised to catalyze the formation of carbamylphosphate [20, 21]. Similarly, determination of intramitochondrial concentration of L-arginine indicated that it was approx. five-fold higher than its NAGS activation constant, suggesting that NAGS molecules are saturated with L-arginine in the mitochondria [22, 23]. Moreover, the addition of glutamine as a source of ammonia to isolated hepatocytes resulted in increased ureagenesis but not elevated NAG levels [20]. Finally, there was no difference in the magnitude of increase of hepatic NAG content and CPS1 activity when mixtures of amino acids with or without arginine were injected into the peritoneum [24]. The first step towards resolving the role of L-arginine in the production of NAG is to determine whether ureagenesis becomes impaired when NAGS cannot bind L-arginine.

The NAGS knockout (*Nags*^−/−^) mouse is an animal model of NAGS deficiency that can survive into adulthood and reproduce when treated with N-carbamylglutamate and L-citrulline (NCG+Cit) from birth, and develops acute hyperammonemia when this treatment is withdrawn. Thus, the *Nags*^−/−^ mouse is uniquely suitable for studying NAGS function *in vivo*. In this study, we tested the effect of a mutant NAGS protein that cannot bind L-arginine, and whether this mutant can restore urea cycle function. To analyze the behavior of *Nags*^−/−^ mice, we employed either a Home Cage Behavioral system or voluntary wheel to quantify their activity during a healthy, treated state, and as they develop hyperammonemia following the withdrawal of NCG+Cit. Then, to assess the phenotype of NAGS E354A, we used gene transfer mediated by the adeno associated virus (AAV) to deliver either wild-type or mutant NAGS to *Nags*^−/−^ mice, and evaluated survival and plasma ammonia following NCG+Cit withdrawal. Finally, we describe the biological consequences of a patient with *NAGS* deficiency associated with a mutant *NAGS* gene, which encodes a protein that cannot bind L-arginine.

## Results

### Non-invasive tracking of hyperammonemia

We used either the Home Cage Scan behavioral monitoring system or voluntary wheel to chronologically assess behavioral and activity changes that occur during the development of hyperammonemia in the *Nags*^−/−^ mice. Six behaviors were monitored with the Home Cage Scan system: eating, drinking, hanging, walking, grooming, and rearing up; we also determined whether time to cessation of each activity differed depending on the withdrawal of the NCG+Cit treatment at either 10 AM or 4 PM. Mouse activity was recorded for 48 hours.

During the first 24-hour period mice were treated with NCG+Cit, there was no significant difference between *Nags*^−/−^ and wild-type mice in any measured behaviors (Figure S1A). Following NCG+Cit withdrawal at 10 AM, the *Nags*^−/−^ mice started to cease drinking, hanging, eating, rearing up, grooming and walking between 4.5 and 17.5 hours (Figures 1A and S1B). Drinking and hanging were the first two activities to cease, followed by eating and rearing up; grooming and walking continued until 17-24 hours after withdrawal of NCG+Cit (Figures 1A and S1B). In contrast, wild type mice appeared to exhibit the same activity levels compared to the first 24-hour period indicating that NCG and L-citrulline have no effect on the activity of healthy mice (Figure S1C).

**Figure 1.**
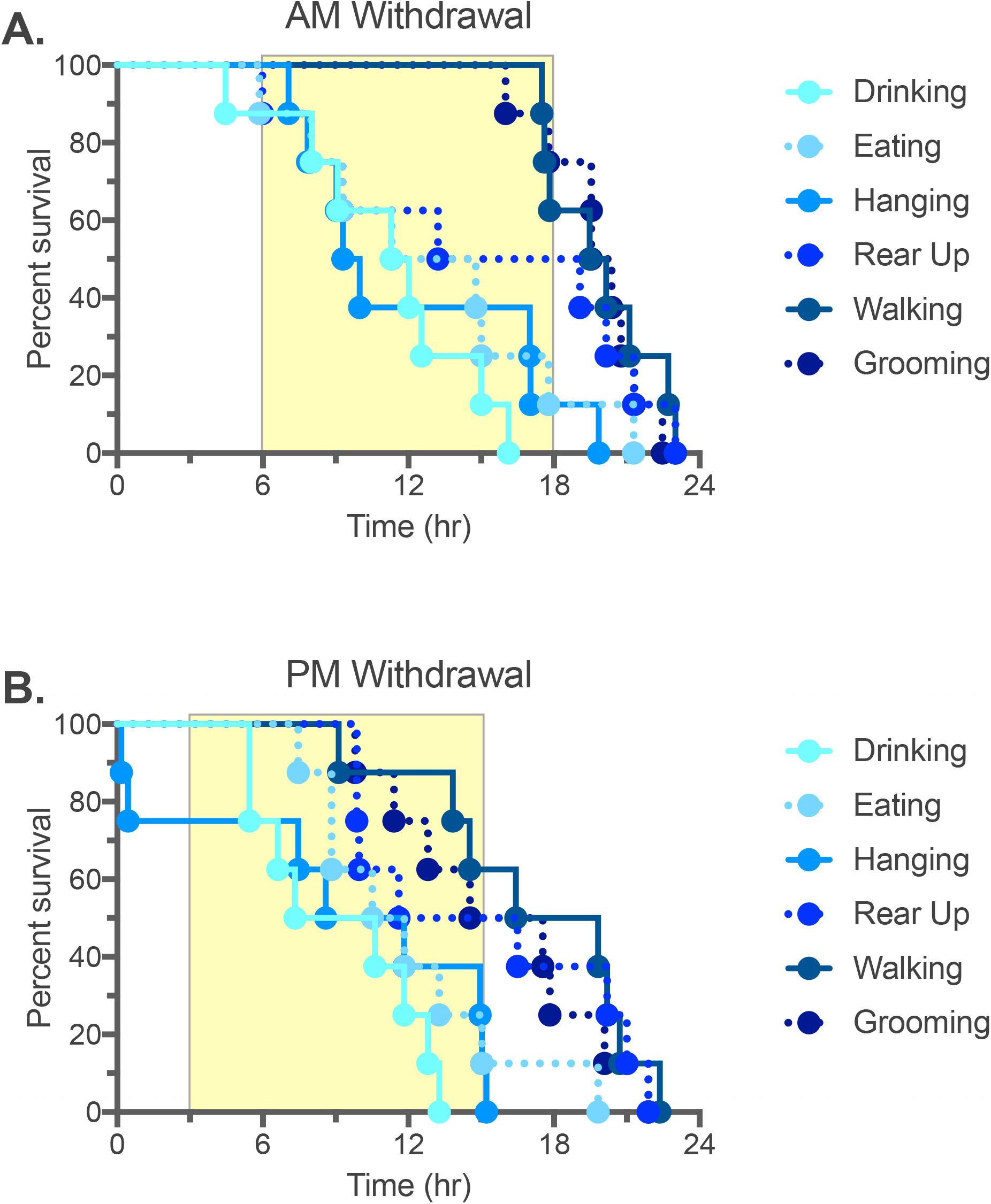
Behavioral changes of the Nags^−/−^ mice. Cessation of six behaviors as *Nags*^−/−^ mice develop hyperammonemia upon withdrawal of NCG+Cit supplementation at 10 AM (**A**) and 4 PM (**B**). Shaded rectangles indicate dark periods.

We thought that the timing of NCG+Cit withdrawal might affect the timing of symptoms related to hyperammonemia because mice are not active during the daytime, and possibly do not drink a significant amount of water during the light compared to dark part of the cycle. To account for circadian rhythm effects, the experiment was repeated with the NCG+Cit withdrawal at 4 PM, just prior to the active period. As in the previous experiment, the behaviors of the *Nags*^−/−^ and wild-type mice were similar before withdrawal of the NCG+Cit treatment (Figure S2A). The *Nags*^−/−^ mice started to cease drinking, eating and hanging behaviors between 10 min. and 5 hours after evening withdrawal of NCG+Cit treatment (Figures 1B and S2B). The cessation of rearing up was similar when NCG+Cit were withdrawn at either 10 AM or 4PM from the *Nags*^−/−^ mice, while they stopped walking and grooming between 9 and 22 hours after withdrawal of the NCG+Cit treatment at 4PM (Figures 1B and S2B). The behavior of the wild-type mice was similar before and after the evening withdrawal of the NCG+Cit treatment (Figure S2C). Although the Home Cage Scan system provides detailed information about multiple mouse behaviors, it has low throughput and we sought to develop a higher throughput method for evaluation of the *Nags*^−/−^ behavior after withdrawal of the rescue chemicals.

### Rescue of Nags^−/−^ phenotype with AAV gene therapy

NCG is an FDA approved drug for *NAGS* deficient patients; however, the life-long therapy is expensive. A more efficient treatment would be transfer of the *NAGS* coding sequence into hepatocytes. Pre-clinical studies have demonstrated the success of correcting ornithine transcarbamylase deficiency in mice using AAV gene therapy [25]. Therefore, we evaluated whether AAV-based gene transfer of the mouse NAGS (mNAGS) can rescue *Nags*^−/−^ mice from hyperammonemia after withdrawal of the NCG treatment, and whether the *Nags* gene promoter can be used to control expression of the delivered *Nags* gene. The number of AAV viral particles needed for the rescue of *Nags*^−/−^ mice and the corresponding expression levels of mNAGS mRNA and protein were determined first. Expression of mNAGS was controlled either by the human thyroxine-binding-globulin (TBG) or natural mouse *Nags* promoter. To determine the dose of vector needed for the rescue of *Nags*^−/−^ mice we measured the duration of activity of *Nags*^−/−^ mice on the voluntary wheel after they were injected with 10^9^, 10^10^, or 10^11^ of either AAV2/8.TBG.mNAGS or AAV2/8.NAGS.mNAGS vector. Control mice were injected with 10^11^ particles of the AAV2/8TBG vector without the mNAGS coding sequence insert (null vector). Five days after injection, mice were transferred into cages with voluntary wheels and, after an acclimation period, their activity on the wheel was monitored for 48 hours before and after withdrawal of NCG from the drinking water. L-citrulline supplementation continued throughout the experiment to ensure endogenous biosynthesis of arginine [26]. All *Nags*^−/−^ mice injected with 10^10^ and 10^11^ particles of either AAV2/8TBG.mNAGS or AAV2/8NAGS.mNAGS appeared healthy and were active for 48 hours after withdrawal of NCG from the drinking water indicating complete rescue by these two doses of viral particles (Figures 2A and 2B). The lowest dose of 10^9^ viral particles of either vector did not rescue *Nags*^−/−^ mice although some did not develop hyperammonemia and cease activity on the wheel close to 48 hours after withdrawal of NCG (Figures 2A and 2B).

**Figure 2.**
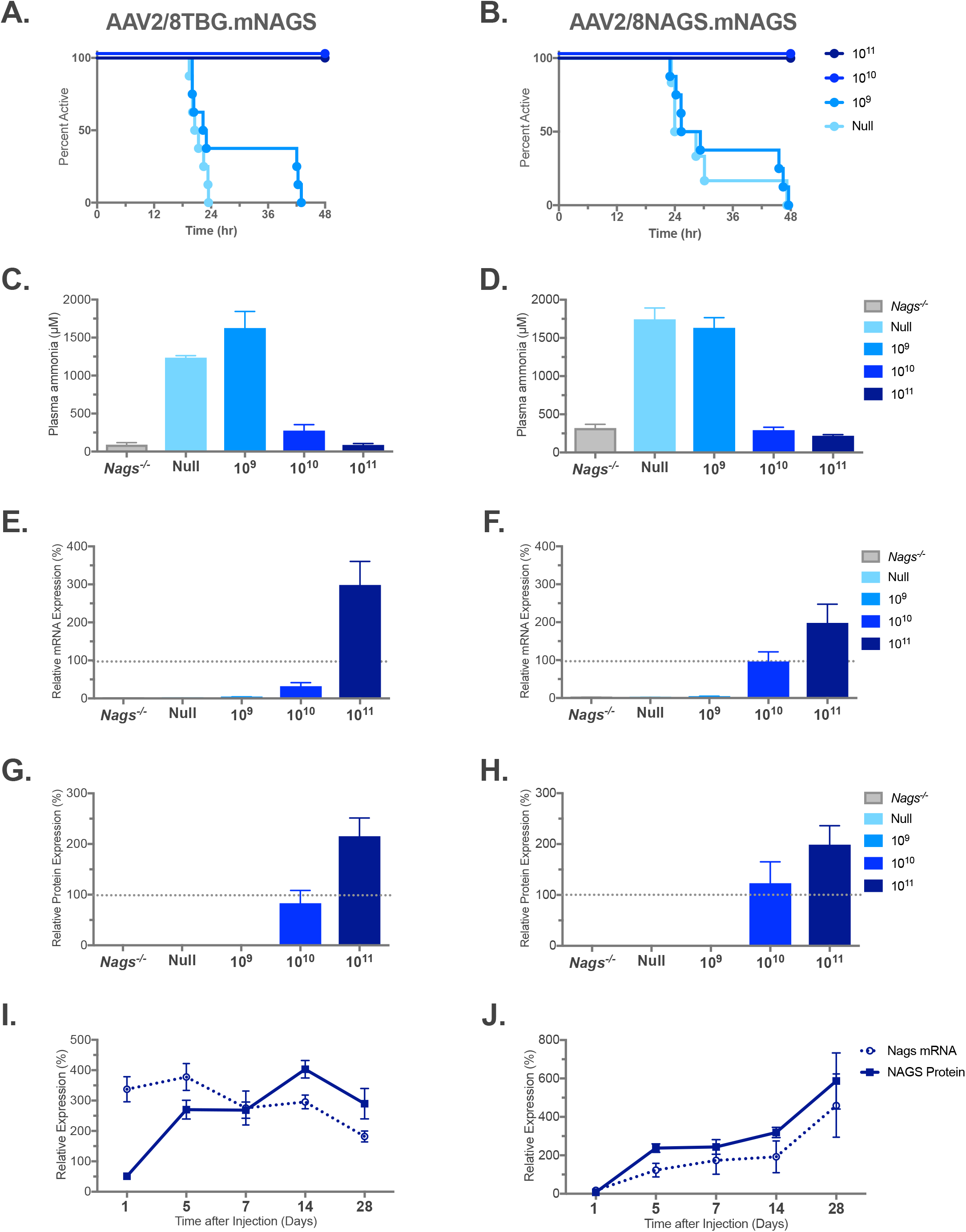
Phenotype of Nags^−/−^ mice after AAV-mediated gene transfer. Duration of activity on the voluntary wheel (**A** and **B**), plasma ammonia concentrations (**C** and **D**), abundance of the liver NAGS mRNA (**E** and **F**) and protein (**G** and **H**) after withdrawal of NCG. Abundance of the NAGS mRNA and protein in the livers of *Nags*^−/−^ mice after single injection of 10^11^ viral particles of either AAB2/8TBG.mNAGS (**I**) or AAB2/8NAGS/mNAGS (**J**).

Plasma ammonia and the expression of mNAGS mRNA and protein were measured in mice that stopped running on the wheel and at the end of the 48-hour activity monitoring period in the mice that were rescued by gene transfer. Plasma ammonia in *Nags*^−/−^ mice supplemented with NCG+Cit was used as control. Mice injected with the null vector and mice injected with 10^9^ particles of either AAV2/8TBG.mNAGS or AAV2/8NAGS.mNAGS developed severe hyperammonemia after they stopped running on the wheel while plasma ammonia was similar to baseline in mice injected with 10^10^ and 10^11^ doses of either vector (Figures 2C and 2D). Real-time PCR was used to determine the abundance of the mNAGS mRNA and immunoblotting was used to determine the abundance of the mNAGS protein. The *Nags*^−/−^ mice supplemented with NCG+Cit and wild-type mice were used as negative and positive controls for mRNA and protein measurements. The mNAGS mRNA and protein were undetectable in the livers of *Nags*^−/−^ mice injected with the null vector (Figures 2E-2H). The abundance of NAGS mRNA was below 10% of the wild-type levels while NAGS protein was undetectable in mice injected with 10^9^ particles of either AAV2/8TBG.mNAGS or AAV2/8NAGS.mNAGS vector (Figures 2E-2H and S3). The average abundance of mNAGS mRNA in mice injected with 10^10^ AAV2/8TBG.mNAGS particles was below wild-type level (32±9% of wild-type, mean±SEM, n=8; Figure 2E) but the abundance of mNAGS protein in these mice was similar to wild-type levels (83±25% of wild-type, mean±SEM, n=8; Figures 2G and S3). The average abundance of mNAGS mRNA and protein in *Nags*^−/−^ mice injected with 10^11^ particles of the AAV2/8TBG.mNAGS vector were 298±62% and 215±36% (mean±SEM, n=8) of the mRNA and protein levels in wild-type mice, respectively (Figures 2E, 2G and S3). In the *Nags*^−/−^ mice injected with 10^10^ AAV2/8NAGS.mNAGS particles average mNAGS mRNA and protein levels were similar to the abundance of mNAGS mRNA and protein in the wild-type mice, 96±25% and 123±42%, respectively(mean±SEM, n=8) while *Nags*^−/−^ mice injected with the highest dose of the AAV2/8NAGS.mNAGS vector had about two-fold higher levels of mNAGS mRNA and protein, 199±49% and 199±37% (mean±SEM, n=8), respectively, than wild-type mice (Figures 2F, 2H and S3).

We also determined the abundance of mNAGS mRNA and protein 1, 5, 7, 14 and 28 days after a single injection of 10^11^ particles of either AAV2/8TBG.mNAGS or AAV2/8NAGS.mNAGS vector. In mice injected with AAV2/8TBG.mNAGS vector mNAGS mRNA and protein were expressed one day after injection (Figure 2I). The abundance of mNAGS mRNA peaked on the fifth day after injection and decreased thereafter while mNAGS protein abundance peaked on day 14 and was lower 28 days after injection (Figure 2I). Expression patterns of mNAGS mRNA and proteins were different in mice injected with the AAV2/8NAGS.mNAGS vector. In these mice average abundance of NAGS mRNA and protein was less than 20% of the NAGS mRNA and protein abundance in wild-type mice one day after injection of AAV2/8-NAGS.mNAGS and increased thereafter (Figure 2J).

### Functional testing of arginine-insensitive NAGS in vivo

To determine the contribution of L-arginine to NAGS function, we used the above AAV-based gene transfer and activity measurements on the voluntary wheel to determine whether the E354A arginine-insensitive mutant mNAGS [27] can rescue *Nags*^−/−^ mice from hyperammonemia. The *Nags*^−/−^ mice were injected with either 10^10^ or 10^11^ viral particles of the AAV2/8NAGS.mNAGS-E354A vector. Five days after injection the mice were placed in cages with a voluntary wheel, allowed two days to acclimate to the new environment and their activity was recorded for 48 hours before and after withdrawal of NCG from the drinking water. Mice injected with 10^11^ viral particles of either AAV2/8NAGS.mNAGS (wild type) or AAV2/8TBGnull were a positive and negative control, respectively. The mice that received either AAV2/8NAGS.mNAGS-E354A or AAV2/8NAGS.mNAGS remained active for 48 hours after withdrawal of NCG, while the activity of mice injected with the null vector ceased between 10 and 45 hours post NCG withdrawal (Figure 3A). Although mice that received the E354A mutant NAGS appeared healthy and remained active for 48 hours after withdrawal of NCG, their plasma ammonia was elevated with inverse correlation to the viral vector dose (Figure 3B) and there was a trend toward higher plasma glutamine levels in mice that received 10^10^ viral particles of the AAV2/8NAGS.mNAGS-E354A vector as compared to the 10^11^ dose (Figure 3C). Measurements of the mRNA and protein abundance revealed no difference in expression levels between the wild-type and E354A mutant mNAGS in the livers of *Nags*^−/−^ mice that were injected with the same dose of either AAV2/8NAGS.mNAGS or AAV2/8NAGS.mNAGS-E354A (Figures 3D, 3E and S4). This suggests that E354A mutant NAGS cannot produce sufficient amount of NAG needed for conversion of all nitrogenous waste into urea.

**Figure 3.**
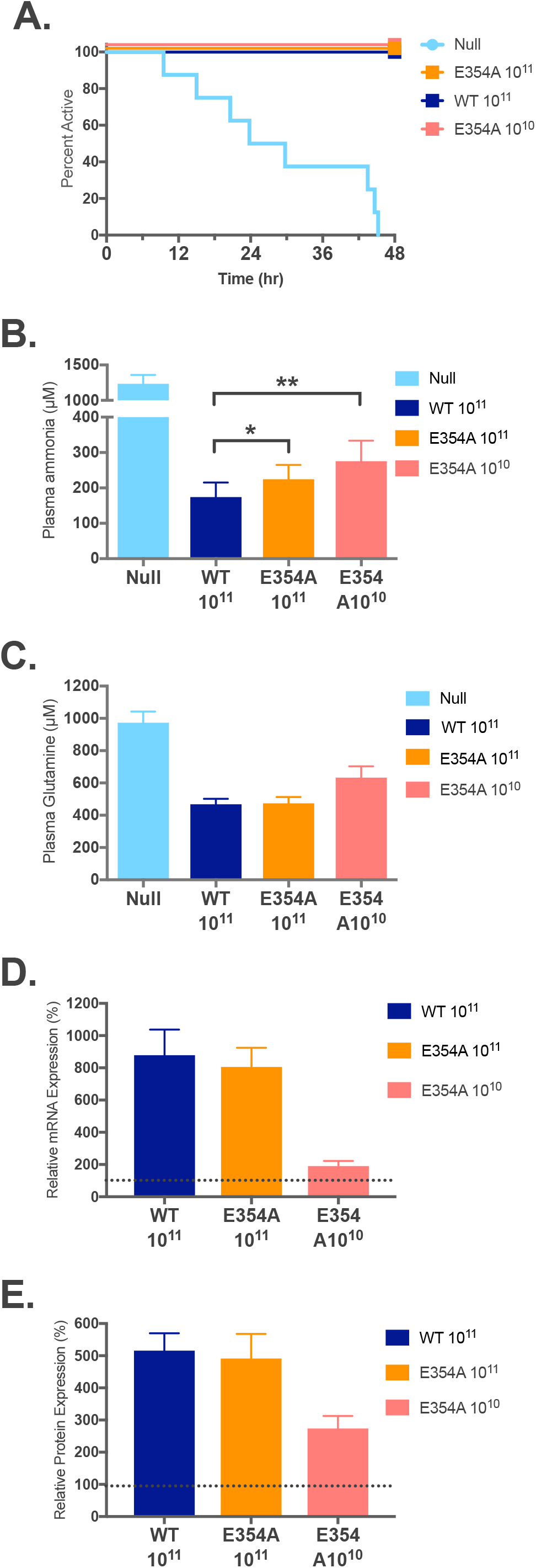
Phenotype of Nags^−/−^ mice that received AAV2/8NAGS.mNAGS-E354A gene therapy. Duration of activity on the voluntary wheel (**A**), plasma ammonia (**B**) and glutamine (**C**) concentrations, abundance of the liver NAGS mRNA (**D**) and protein (**E**) after withdrawal of NCG.

We analyzed activity of the *Nags*^−/−^ mice that were injected with 10^11^ viral particles of either AAV2/8NAGS.mNAGS or AAV2/8NAGS.mNAGS-E354A to determine whether hyperammonemia affects their behavior on the voluntary wheel. The 48 hours before and after withdrawal of NCG supplementation were divided into 6-hour epochs followed by calculation of the running time and distance for each mouse during each epoch. Before withdrawal of NCG supplementation, both NCG and NAG should be present in the livers to activate CPS1, while after withdrawal of NCG, CPS1 should be activated by the NAG produced by either E354A or wild-type NAGS that were introduced by gene transfer. Running time of the *Nags*^−/−^ mice that received either wild-type or E354A mNAGS did not differ before withdrawal of the NCG supplementation (Figure 4A); after withdrawal of NCG there was a trend (p=0.06) towards longer running time for mice that received E354A mNAGS (Figure 4B). Running distance was similar for mice that received either wild-type or E354A mNAGS before withdrawal of NCG, while after NCG withdrawal the distance run by the mice that received the E354A mNAGS was longer than the distance run by the mice that received wild-type mNAGS (Figure 4D) suggesting that mice may run faster when they are hyperammonemic. To confirm this, we determined the length of each running session on the wheel, the distance a mouse run during that session and its velocity for the 6 hour epochs after withdrawal of NCG supplementation. At nighttime *Nags*^−/−^ mice that received E354A mutant mNAGS gene run longer distance because during running sessions they run faster than mice that received wild-type mNAGS gene (Figure 4E).

**Figure 4.**
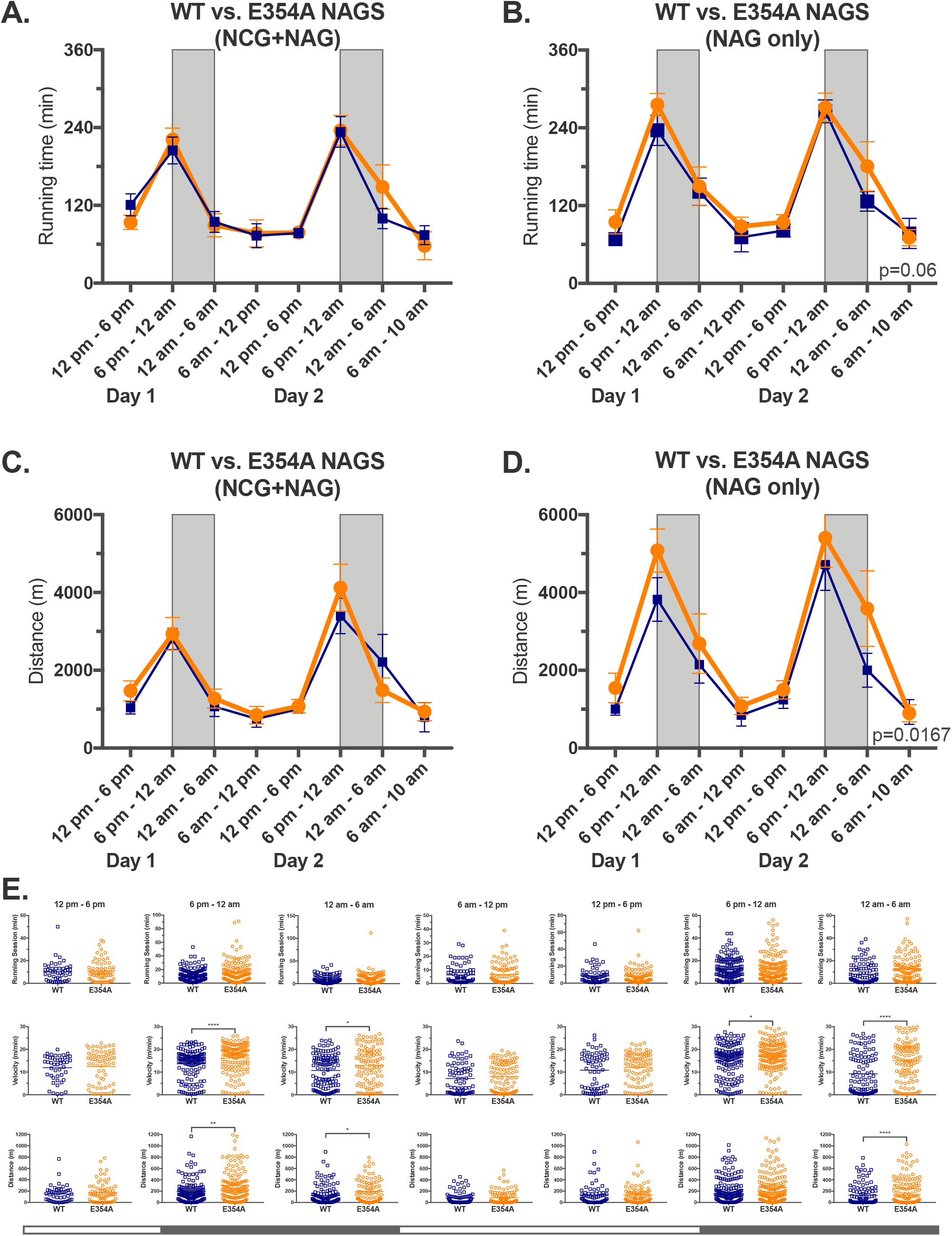
Behavior of Nags^−/−^ mice that received the same dose of either AAV2/8NAGS.mNAGS-E354A or AAV2/8NAGS.mNAGS gene therapy. **A.** Average running time during 6 hr. epochs before withdrawal of NCG supplementation. **B.** Average running time during 6 hr. epochs after withdrawal of NCG supplementation. **C.** Average running distance during 6 hr. epochs before withdrawal of NCG supplementation. **D.** Average running distance during 6 hr. epochs after withdrawal of NCG supplementation. **E.** Length of running sessions, velocity and distance during each running session in 6 hr. epochs after withdrawal of NCG supplementation. Orange – AAV2/8NAGS.mNAGS-E354A. Blue – AAV2/8NAGS.mNAGS. Shaded rectangles and bars indicate nighttime when mice are more active. * – p*<*0.01, ** – p*<*0.001, **** – p*<*0.0001.

### NAGS deficiency due to arginine-insensitive NAGS

The clinical relevance of arginine-insensitive NAGS was confirmed when a patient with late-onset *NAGS* deficiency was found to be a compound heterozygous for two sequence variants, NM 153006.2:c.1080G*>*T (NP 694551.1:p.E360D) and NM 153006.2:c.426+3G*>*C, while we were testing the function of the arginine insensitive E354A mutant mNAGS in mice. Functional testing of the c.426+3G*>*C sequence variant indicated that it could diminish NAGS expression, presumably due to impaired mRNA splicing (Figure S5). The p.E360D amino acid substitution affects a glutamate residue in human NAGS (hNAGS) that corresponds to the E354 in mNAGS [27, 28]. To determine the functional consequences of this substitution, the E360D and E354D mutations were engineered into hNAGS and mNAGS, respectively, followed by overexpression in *E. coli* and purification of mutant and wild-type recombinant hNAGS and mNAGS. We tested whether the enzymatic activity of the four recombinant proteins increases in the presence of L-arginine and whether they can bind L-arginine. The results revealed that the enzymatic activities of the wild-type hNAGS and mNAGS doubled in the presence of 1 mM L-arginine, while the activities of the E360D hNAGS and E354D mNAGS did not change in the presence of 1 mM L-arginine (Table 1). Thermofluor assay was used to determine the effect of L-arginine on thermal unfolding of wild-type and mutant recombinant hNAGS and mNAGS; D-arginine, which does not bind to mNAGS [29], was used as a control. The Tm (temperature at which half of the protein molecules are unfolded), of both hNAGS and mNAGS increased by 4.60±0.05°C and 4.70±0.12°C (mean±SD, n=3), respectively, in the presence of L-arginine (Figures 5A and 5B), while the Tm of the E360D hNAGS and E354D mNAGS remained the same in the presence of either L- or D-arginine (Figures 5C and 5D). This suggests that these mutant hNAGS and mNAGS proteins cannot bind L-arginine, explaining the absence of its effect on NAGS enzymatic activity.

**Figure 5.**
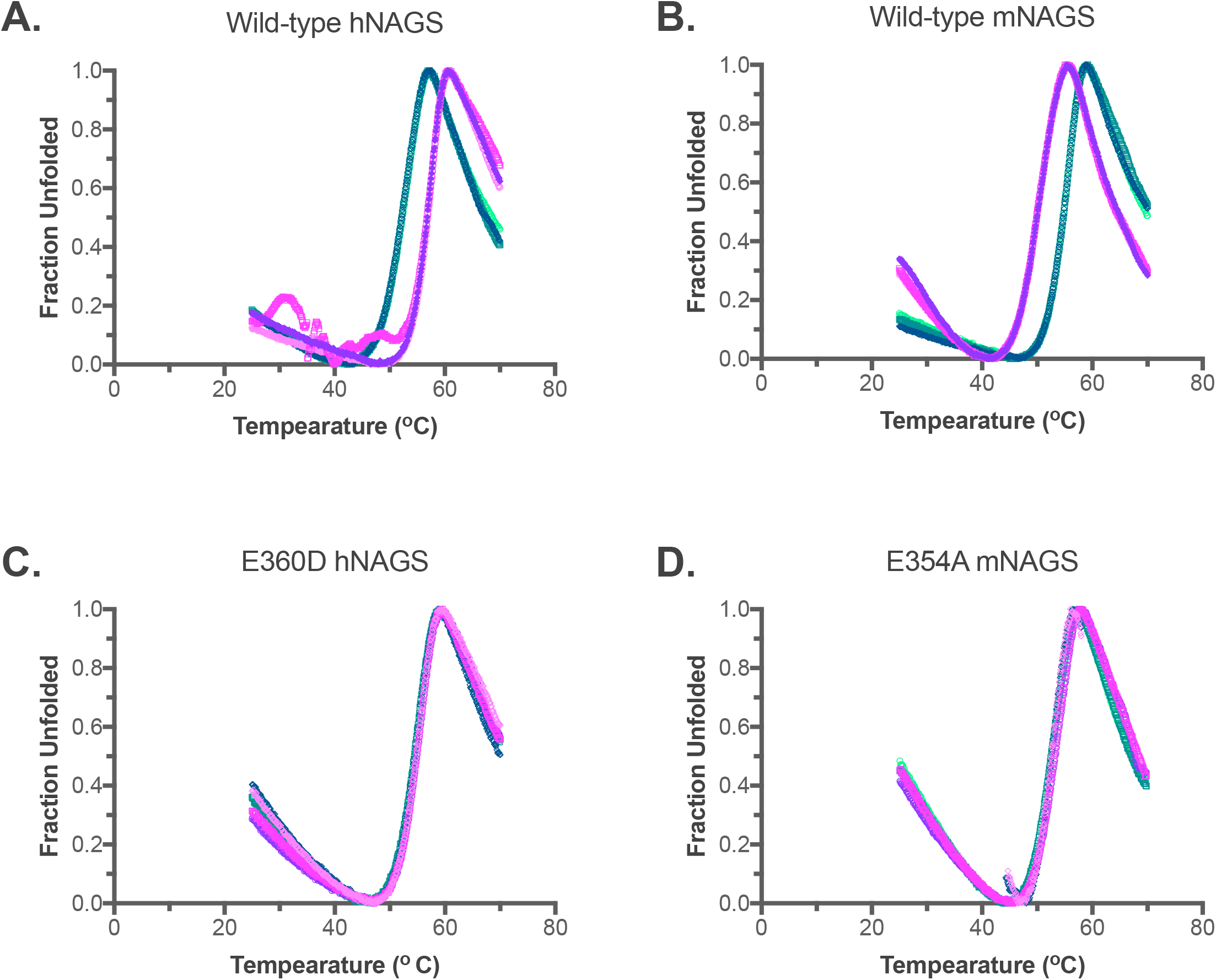
Thermofluor analysis. Wild-type human (**A**) and mouse (**B**) NAGS, E360D human NAGS (**C**), and E354D mouse NAGS (**D**) in the presence of either L-arginine (purple) or D-arginine (teal).

**Table 1.** Effect of L-arginine on enzymatic activities of either wild-type or mutant human and mouse NAGS.

| Protein | NAGS Specific Activity<br>($\mu\text{moles min}^{-1}\text{mg}^{-1}$) | |
| --- | --- | --- |
|  |  | + 1 mM L-Arg |
| WT hNAGS | 7.17±1.02<sup>a</sup> | 13.98±1.95 |
| E360D hNAGS | 10.15±0.10 | 10.05±0.20 |
| WT hNAGS | 13.29±0.08 | 23.94±0.37 |
| E354D mNAGS | 8.28±0.13 | 8.24±0.06 |

<sup>a</sup>Enzymatic activities are averages of three measurements and associated standard errors.

| **Protein** | **NAGS Specific Activity**<br>($\mu\text{moles min}^{-1}\text{mg}^{-1}$) | |
| --- | --- | --- |
|  |  | **+ 1 mM L-Arg** |
| Wild-type human NAGS | 7.17±1.02<sup>a</sup> | 13.98±1.95 |
| E360D human NAGS | 10.15±0.10 | 10.05±0.20 |
| Wild-type mouse NAGS | 13.29±0.08 | 23.94±0.37 |
| E354D mouse NAGS | 8.28±0.13 | 8.24±0.06 |

<sup>a</sup>Enzymatic activities are averages of three measurements and associated standard errors.

## Discussion

The results presented herein provide direct evidence that binding of L-arginine to mammalian NAGS is required for its normal function. The *Nags*^−/−^ mice that express the E354A mutant mNAGS, which is not activated by L-arginine [27], are hyperammonemic despite similar expression levels of the mutant and wild-type mNAGS mRNA and protein (Figure 3). Moreover, we describe a case of a hyperammonemic patient found to have a E360D substitution in NAGS; this mutated protein neither binds nor is activated by L-arginine, suggesting that inability to bind L-arginine impairs the function of human NAGS, leading to decreased ureagenesis and hyper-ammonemia. Another possibility is that inability to bind L-arginine has a dominant negative effect on the function of NAGS heterotetramers that consist of wild-type and mutant subunits in the patient with NAGS deficiency.

The voluntary wheel was used to elucidate the biological consequences arginine-insensitive NAGS on urea cycle function and hyperammonemia development. The hyperammonemic *Nags*^−/−^ mice that express the E354A mutant NAGS and the *Nags*^−/−^ mice that expressed wild-type NAGS had different patterns of activity on the voluntary wheel. Hyperammonemic mice, that received E354A mutant NAGS, run longer distance on the voluntary wheel during nighttime than the mice that received the same dose of wild type NAGS. This was because hyperammonemic mice run faster than the mice that received wild-type NAGS since the length of running session did not differ between the two groups of mice. This suggests that hyperammonemic mice may be hyperactive, similar to patients with urea cycle disorders who can have symptoms of attention deficit hyperactivity disorder [30, 31].

In mammals and other animals that detoxify ammonia via the urea cycle, the function of NAGS is to provide a cofactor for either CPS1 or CPS3. However, in microbes and plants, NAGS catalyzes the first reaction of L-arginine biosynthesis [32-36]. The effect of L-arginine on NAGS enzymatic activity changes during evolution as well. L-arginine acts as a negative feedback signal in organisms with Larginine biosynthesis, while it partially inhibits NAGS activity in fish and activates NAGS in mammals [27]. Despite these differences, the binding site for L-arginine remains conserved in all NAGS enzymes [27, 37]. While feedback inhibition of microbial and plant NAGS proteins provides obvious benefits of preventing unnecessary consumption of precursors and energy when exogenous L-arginine is abundant, the benefits of binding and activation of mammalian NAGS by L-arginine are less clear. One possibility is that activation of mammalian NAGS by L-arginine together with activation of CPS1 by NAG creates a double positive feedback loop that enables sensitive and robust regulation of ammonia detoxification via ureagenesis and effective protection of the central nervous system from ammonia toxicity [27, 38]. Alternatively, L-arginine and its binding to mammalian NAGS could be part of a nitrogen load sensing mechanism that adjusts abundance of urea cycle enzymes in response to increased protein catabolism due to either high dietary protein intake or degradation of cellular proteins. We will be able to test these hypotheses by combining biochemical testing with monitoring of activity and behavior of *Nags*^−/−^mice either on the voluntary wheel or in the Home Cage system.

## Materials and Methods

### Animal husbandry and genotyping

Animals were housed at Children’s National Medical Center and the Veterans Affairs Animal Research Facility and all protocols were approved by The Children’s National Medical Center and V.A. IACUC, Washington, DC. Mice were checked daily for signs of distress and were maintained on a 12:12-h light–dark cycle in a low-stress environment (22°C, 50% humidity and low noise) and given food and water *ad libitum*.

The maintenance of the *Nags*^−/−^ mouse colony has been previously described [39]. For the purposes of these studies, wild type mice were given NCG+Cit supplementation for a 2-week period before the experiments were conducted.

Genomic DNA was isolated either from tail or ear punch biopsy using Gentra Puregene Tissue kit (Qiagen, Valencia, CA) according to manufacturer’s instructions. Mice were genotyped either as described previously [39] or using multiplex PCR with primers Exon6F: 5’-CAG CTT TAG GAG GAC AGG AGA G-3’, Exon6R: 5’-CTT AGA GAC ACA GAC CAG GAG TTA G-3’, NEO-F: 5’-TGC TCC TGC CGA GAA AGT ATC CAT CAT GGC-3’ and NEO-R: 5’-CGC CAA GCT CTT CAG CAA TAT CAC GGG TAG-3’ and the following amplification conditions 2 min. initial denaturation at 95°, followed by 25 cycles of 15 sec. denaturation at 95°C, 20 sec. annealing at 60°C, 45 sec. extension at 72°C, and 5 min. final extension at 72°C. Primers Exon6F and Exon6R amplify 510 bp long wild-type allele while primers NEO-F and NEO-R amplify 340 bp long knockout allele. Genotype of each mouse used in this study was determined before experiments and confirmed at the end of each experiment.

### Home cage monitoring

We used the Home Cage Scan system from CleverSys [13-17] consisting of 4 cameras simultaneously monitoring 4 mice individually housed in separate cages on a 12:12 light: dark cycle. The cage environment was lit by white lights from 6 AM to 6 PM and infrared lights from 6 PM to 6 AM. The computer software allows the researcher to calibrate the cage, providing software with information about the cage environment such as the top/bottom of cage, height of bedding, placement of drinking spout, and food bin areas. The software then uses this information to recognize the animal and behavior with duration greater than 6 frames per second.

Eight 6-weeks-old *Nags*^−/−^ and eight aged-matched wild-type controls were used for this study. Mice were housed with minimal bedding to minimize mounding, which can obscure the mouse during recording. Mice were acclimated to their cages and the environment for 48 hours prior to the study and were recorded for 23 hours beginning at either 10:00 AM or 4 PM. Mice remained on NCG+Cit supplemented water for the first 23 hours. After the first 23 hours, the NCG+Cit supplemented water was replaced with drug-free water before the next 23 hours of recording. Recording of behavior lasted for 23 hours in order to give the researcher time to change the water and recalibrate the cages so that day 2 recording could also start at either 10:00 AM or 4 PM. Random bouts of movies were inspected after the experiment to determine the accuracy of the behaviors called by the recognition software. Incorrect behaviors were manually corrected by the researcher. Results are presented as the percent of animals displaying a behavior.

The information provided by the software provides the researchers with more than 30 different behaviors. We chose to group some of these behaviors into six categories for data analysis (Table S1). The software recognizes and distinguishes the behaviors rear up, come down from partially reared, come down to partially reared, rear up from partially reared, rear up to partially reared, remain reared up, and remain partially reared, which we grouped as “Rear up”. “Hanging” regroups the behaviors hang cuddled, hang vertical from hang cuddled, remain hang vertical, and remain hang cuddled as defined by the software. “Eat” regroups the software distinguished behaviors eat and chew. “Walk” regroups the software distinguished behaviors walk left, walk right, and walk slowly (Table S1).

### Measurements of mouse activity with voluntary wheels

Eight 6-8-weeks-old mice per experimental group were kept individually in cages equipped with a running wheel with the magnet (Minimitter Company, Inc. Surriver or Vers 4.1). Running wheel system was monitored by online computer using Vital View data acquisition system (StarrLife, Inc). The number of turns of each wheel was recorded in 1 min. bins. The running distance and velocity were calculated based on a 0.36 m wheel circumference. After transfer from their home cages to cages with voluntary wheels, mice were allowed 48 hr. acclimation to the new environment. Following acclimation period, activity of mice supplemented with NCG+Cit in the drinking water was recorded for 48 hours; NCG was removed from the drinking water on the third day at 12 PM and activity was recorded for additional 48 hours. Mice that became inactive due to hyperammonemia were euthanized and their plasma and livers were collected and stored at −80°C for molecular and biochemical analyses. Physical activity was measured as running time, distance, and velocity.

### Plasmid construction and AAV vector preparation

The 1593bp mNAGS coding sequence identical to GenBank ID AF462069.1 was synthesized by DNA2.0, Inc. The AAV2/8.TBG.mNAGS plasmid was generated by inserting mNAGS coding sequence into AAV2/8.TBG plasmid [25] using restriction endonucleases *Not*I and *Bgl*II. Correct sequence between AAV inverted repeats of the AAV2/8.TBG.mNAGS construct was verified by DNA sequencing. For the AAV2/8.NAGS.mNAGS construct the TBG promoter was replaced with the mouse *Nags* promoter [40]. Mouse genomic DNA was extracted following the standard protocol using DNeasy Blood and Tissue Kit, Qiagen (Valencia, CA). The 684 bp fragment harboring mouse *Nags* promoter was amplified using primers 5’-CAG ATC CGG CGC GCC CTC TCT ATA ATA TGT AGC CCC-3’ and reverse primer 5’-GAC CAA CTT CTG CAG GAC GAC AAC CAA ACCC ACT CG-3’, and the following PCR conditions: 5 min. initial denaturation at 95°C followed by 30 cycles of 30 sec. denaturation at 95°C, 30 sec. annealing at 62.5°C, 1 min. extension at 72°C, and 7 min. final extension at 72°C. The TBG promoter of the AAV2/8.TBG.mNAGS plasmid was replaced with the amplification product using In-Fusion Cloning Kit (Clontech) according to manufacturer’s instructions. Correct sequence between AAV inverted repeats of the AAV2/8.NAGS.mNAGS construct was verified by DNA sequencing. The E354A mutation was introduced using 5’-GCA CGC TGC TCA CGG CAC TCT TTA GTA ACA AGG GC-3’ mutagenic primer and the QuikChange Lightning Site-directed mutagenesis kit (Agilent) according to manufacturer’s instruction. Correct sequence between AAV inverted repeats of the resulting AAV2/8.NAGS.mNAGS-E354A construct was verified by DNA sequencing. QIAGEN Plasmid Giga Kit was used to purify approx. 2.4 *µ*g/l of AAV2/8.TBG.mNAGS, AAV2/8.NAGS.mNAGS and AAV2/8NAGS.mNAGS-E354A plasmids. AAV particles containing each of these three plasmids were prepared by the Penn Vector Core, Gene Therapy Program, University of Pennsylvania School of Medicine, USA.

### Delivery of mNAGS into *Nags*^−/−^ mice

AAV viral particles were stored as suspension in sterile phosphate buffered saline (PBS) with 5% glycerol at −80°C. The viral particle suspension was thawed, diluted to concentrations indicated in the text, and delivered via tail vein injections of 100 *µ*l particle suspension. After the injection experimental mice were kept in the home cages for five days prior to voluntary wheel experiments. In mRNA and protein persistence experiments, mice were kept in home cages after injections of the AAV viral particles and were supplemented with NCG+Cit during this time.

### Plasma ammonia and amino acid analysis

Blood was collected from euthanized mice by cardiac puncture. Blood was immediately centrifuged at 7,500 x g for 5 min. at 4°C, and the plasma was stored at −80°C until further analysis. The Ammonia Assay Kit (Abcam, Cambridge, MA) was used to determine plasma ammonia according to the manufacturer’s protocol.

Plasma amino acid concentrations were measured using ion-exchange chromatography on a High-Speed Amino Acid Analyzer L-8800/L-8800A (Hitachi). Plasma proteins were precipitated with an equal volume of 7% sulfosalicylic acid and centrifuged for 10 min. at 13000 rpm. 5 *µ*l of 2.5 N LiOH and 10 *µ*l stock internal standard (S-2-aminoethyl-L-cysteine hydrochloride) were then added to 100 *µ*l of plasma supernatant before loading samples into the analyzer. Amino acids were quantified using L-8800 ASM software package.

### RNA extraction and real-time PCR

RNA extraction was performed as per the manufacturer’s instruction using RNeasy Mini Kit (Qiagen, USA). A total of 2 *µ*g of RNA were transcribed into cDNA using the High-Capacity cDNA reverse Transcription kit (Applied Biosystems, USA) according to the manufacturer’s instructions. For the real time PCR analysis, 1 *µ*l of mNAGS gene probe Mm00467530 m1 (TaqMan), 10 *µ*l of TaqMan Gene Expression Master Mix and 4 *µ*l of cDNA were combined in 20 *µ*l reaction and subjected to PCR using AB 7900 Real-Time PCR System. Abundance of mNAGS mRNA was determined using ΔΔCt method [41]; 18S rRNA gene was used for normalization.

### Immunoblotting

Frozen liver samples were pulverized with liquid nitrogen and the cells were lysed using lysis buffer (50 mM Tris pH 7.5, 1 mM EDTA, 1% NP40, 250 mM Sucrose, 0.1% SDS and Protease Inhibitor Cocktail tablets, Roche). Protein quantification was performed using the Quick StartTM Bradford Protein Assay (Bio-Rad) according to manufacturer’s instructions. 50 *µ*g of proteins from the liver lysate were resolved using 4-20% gradient polyacrylamide gel (Bio-Rad, USA) and transferred to nitrocellulose membrane using Trans-Blot Turbo Transfer System (Bio-Rad, USA). Membranes were then blocked with the Starting Block (TBS) Blocking buffer (Thermo Scientific, Rockford, IL) with 0.5% Surfact-Amps 20 (Thermo Scientific, Rockford, IL) overnight at 4°C. Blocked membranes were probed with the rabbit anti-mNAGS antibody (1:3000 dilution) for 1 hr. at room temp. Primary antibody was detected with the peroxidase conjugated goat anti-rabbit (1:20,000 dilution) secondary antibody (Thermo Scientific, Rockford, IL) and visualized using Super Signal West Pico Chemiluminescent Substrate (Thermo Scientific, Rockford, IL). The band intensities were quantified using ChemiDocTM Touch Imaging System instrument with Image Lab software, version 5.2.1 (Bio-Rad, USA).

### Plasmid construction, recombinant protein overexpression, purification, and biochemical characterization

Plasmids pET15bhNAGS-M and pET15bmNAGS-M [42] were used for overexpression of mNAGS and hNAGS in *E. coli*. Mutagenic primers 5’-GCA CGC TGC TCA CTG ACC TCT TTA GCA ACA AGG-3’ and 5’-GCA CGC TGC TCA CGG ACC TCT TTA GTA ACA AGG-3’ and were used to generate constructs pETbhNAGS-E360D and pET15bmNAGS-E354D using QuikChange Lightning Site-directed mutagenesis kit (Agilent) according to manufacturer’s instruction. Recombinant NAGS proteins were purified using nickel affinity chromatography as described previously [27, 43]. Quality of purified proteins was assessed using SDS-PAGE.

Specific activities of purified enzymes were measured as described previously [44]. Thermal shift assays were performed in a 96 well plate format using a QuantStudio 7Flex Real-Time PCR System (Applied Biosystems). Protein unfolding was monitored by measuring the change in fluorescence intensity of SYPRO Orange (Invitrogen) while ramping temperature from 4°C to 99°C. Wells contained 1 *µ*g of enzyme in 50 mM potassium phosphate pH 7.5, 300 mM KCl, 20% glycerol, 250 mM imidazole, 10 mM *β*-mercaptoethanol (BME), 0.006% Triton X-100, 1% acetone, 50x SYPRO Orange and 10 mM of either L- or D-arginine.

### Statistical analysis

The Kaplan-Meier estimator was used to analyze the time to cessation of activity on the voluntary wheel. Two-way ANOVA was used to analyze differences in activity on the voluntary Wheel. Mann-Whiney test was used to analyze differences in the length of running sessions, running distance and velocity for during each session. All other data were analyzed using Student’s t-test.

## Author’s contributions

PS generated plasmids that were used for gene transfer, carried out activity measurements on the voluntary wheel, molecular and biochemical analyses of mouse tissues, overexpression, purification and biochemical characterization of recombinant NAGS proteins, and wrote the manuscript. ES carried out analysis of mouse behavior using Home Cage Monitoring System. PU created Python script used for parsing activity data from the voluntary wheel. MM diagnosed patient with NAGS deficiency. MT participated in experimental design and interpretation, and critically reviewed the manuscript. HM participated in experimental design, writing of the manuscript and critically reviewed the manuscript. LC conceived this study, edited and reviewed the manuscript.

## Acknowledgements

This work was supported by Public Health Service Grant R01DK064913 from the National Institutes of Health. AAV gene transfer vectors were prepared by the Penn Vector Core, Gene Therapy Program, University of Pennsylvania School of Medicine, USA.

## Additional Files

Additional file 1 – Supplementary Material

Additional File 1 contains Table S1, Figures S1 - S5, and Supplementary Material and Methods for functional testing of the c.426+3G>C sequence variant.

## References

1. Brusilow SW, Horwich AL: Urea Cycle Enzymes. In: The Metabolic Molecular Bases of Inherited Disease. Edited by Scriver CR, Beaudet AL, Sly WS, Valle D, vol. 2: McGraw-Hill; 2001: 1909–1963.

2. Ah Mew N, Simpson KL, Gropman AL, Lanpher BC, Chapman KA, Summar ML: Urea Cycle Disorders Overview. In: GeneReviews((R)). Edited by Adam MP, Ardinger HH, Pagon RA, Wallace SE, Bean LJH, Stephens K, Amemiya A. Seattle (WA); 1993.

3. Bachmann C, Krahenbuhl S, Colombo JP: Purification and properties of acetyl-CoA:L-glutamate Nacetyltransferase from human liver. Biochem J 1982, 205(1):123–127.

4. Shigesada K, Tatibana M: N-Acetylglutamate synthetase from rat-liver mitochondria. Partial purification and catalytic properties. Eur J Biochem 1978, 84(1):285–291.

5. Sonoda T, Tatibana M: Purification of N-acetyl-L-glutamate synthetase from rat liver mitochondria and substrate and activator specificity of the enzyme. J Biol Chem 1983, 258(16):9839–9844.

6. Grisolia S, Cohen PP: Catalytic role of of glutamate derivatives in citrulline biosynthesis. J Biol Chem 1953, 204(2):753–757.

7. Waterlow JC: The mysteries of nitrogen balance. Nutr Res Rev 1999, 12:25–54.

8. Caldovic L, Morizono H, Gracia Panglao M, Gallegos R, Yu X, Shi D, Malamy MH, Allewell NM, Tuchman M: Cloning and expression of the human N-acetylglutamate synthase gene. Biochem Biophys Res Commun 2002, 299(4):581–586.

9. Caldovic L, Morizono H, Yu X, Thompson M, Shi D, Gallegos R, Allewell NM, Malamy MH, Tuchman M: Identification, cloning and expression of the mouse N-acetylglutamate synthase gene. Biochem J 2002, 364(Pt 3):825–831.

10. Morimoto BH, Brady JF, Atkinson DE: Effect of level of dietary protein on arginine-stimulated citrulline synthesis. Correlation with mitochondrial N-acetylglutamate concentrations. Biochem J 1990, 272(3):671–675.

11. Nissim I, Luhovyy B, Horyn O, Daikhin Y, Nissim I, Yudkoff M: The role of mitochondrially bound arginase in the regulation of urea synthesis: studies with [U-^15^N4]arginine, isolated mitochondria, and perfused rat liver. J Biol Chem 2005, 280(18):17715–17724.

12. Felipo V, Minana MD, Grisolia S: Control of urea synthesis and ammonia utilization in protein deprivation and refeeding. Arch Biochem Biophys 1991, 285(2):351–356.

13. Saheki T, Katsunuma T, Sase M: Regulation of urea synthesis in rat liver. Changes of ornithine and acetylglutamate concentrations in the livers of rats subjected to dietary transitions. J Biochem (Tokyo) 1977, 82(2):551–558.

14. Shigesada K, Tatibana M: Enzymatic synthesis of acetylglutamate by mammalian liver preparations and its stimulation by arginine. Biochem Biophys Res Commun 1971, 44(5):1117–1124.

15. Zollner H: Regulation of the N-acetylglutamate content of rat hepatocytes by the glutamate concentration. Adv Exp Med Biol 1982, 153:197–205.

16. Kawamoto S, Ishida H, Mori M, Tatibana M: Regulation of N-acetylglutamate synthetase in mouse liver. Postprandial changes in sensitivity to activation by arginine. Eur J Biochem 1982, 123(3):637–641.

17. Tuchman M, Holzknecht RA: Human hepatic N-acetylglutamate content and N-acetylglutamate synthase activity. Determination by stable isotope dilution. Biochem J 1990, 271(2):325–329.

18. Beliveau Carey G, Cheung CW, Cohen NS, Brusilow S, Raijman L: Regulation of urea and citrulline synthesis under physiological conditions. Biochem J 1993, 292(Pt 1):241–247.

19. Shigesada K, Tatibana M: Role of acetylglutamate in ureotelism. I. Occurrence and biosynthesis of acetylglutamate in mouse and rat tissues. J Biol Chem 1971, 246(18):5588–5595.

20. Lund P, Wiggins D: Is N-acetylglutamate a short-term regulator of urea synthesis? Biochem J 1984, 218(3):991–994.

21. Van Dijk M, Lund P: N-Acetylglutamate in rat liver during foetal development. Biochem J 1984, 222(3):837–838.

22. Freedland RA, Meijer AJ, Tager JM: Nutritional influences on the distribution of the urea cycle: intermediates in isolated hepatocytes. Fed Proc 1985, 44(8):2453–2457.

23. Horyn O, Luhovyy B, Lazarow A, Daikhin Y, Nissim I, Yudkoff M, Nissim I: Biosynthesis of agmatine in isolated mitochondria and perfused rat liver: studies with ^15^N-labelled arginine. Biochem J 2005, 388(Pt 2):419–425.

24. Stewart PM, Walser M: Short term regulation of ureagenesis. J Biol Chem 1980, 255(11):5270–5280.

25. Wang L, Wang H, Morizono H, Bell P, Jones D, Lin J, McMenamin D, Yu H, Batshaw ML, Wilson JM: Sustained correction of OTC deficiency in spf(ash) mice using optimized self-complementary AAV2/8 vectors. Gene Ther 2012, 19(4):404–410.

26. Windmueller HG, Spaeth AE: Source and fate of circulating citrulline. Am J Physiol 1981, 241:E473–E480.

27. Haskins N, Panglao M, Qu Q, Majumdar H, Cabrera-Luque J, Morizono H, Tuchman M, Caldovic L: Inversion of allosteric effect of arginine on N-acetylglutamate synthase, a molecular marker for evolution of tetrapods. BMC Biochem 2008, 9:24.

28. Caldovic L, Morizono H, Panglao MG, Lopez GY, Shi D, Summar ML, Tuchman M: Late onset N-acetylglutamate synthase deficiency caused by hypomorphic alleles. Hum Mutat 2005, 25(3):293–298.

29. Haskins N, Mumo A, Brown PH, Tuchman M, Morizono H, Caldovic L: Effect of arginine on oligomerization and stability of N-acetylglutamate synthase. Sci Rep 2016, 6:38711.

30. Batshaw ML, Roan Y, Jung AL, Rosenberg LA, Brusilow SW: Cerebral dysfunction in asymptomatic carriers of ornithine transcarbamylase deficiency. N Engl J Med 1980, 302(9):482–485.

31. Kim SH, Lee JS, Lim BC, Kim KJ, Hwang YS, Park JD, Cheon JE, Kim IO, Kim BN, Chae JH: A female carrier of ornithine carbamoyltransferase deficiency masquerading as attention deficit-hyperactivity disorder. Brain Dev 2014, 36(8):734–737.

32. Caldovic L, Tuchman M: N-acetylglutamate and its changing role through evolution. Biochem J 2003, 372(Pt 2):279–290.

33. Terjesen BF, Ronnestad II, Norberg B, Anderson PM: Detection and basic properties of carbamoyl phosphate synthetase III during teleost ontogeny: a case study in the Atlantic halibut (Hippoglossus hippoglossus L.). Comp Biochem Physiol B 2000, 126(4):521–535.

34. Xu Y, Glansdorff N, Labedan B: BioiDetection and basic properties of carbamoyl phosphate synthetase III during teleost ontogeny: a case study in the Atlantic halibut (Hippoglossus hippoglossus L.).ation of N-acetylated intermediates in prokaryotes. BMC Genomics 2006, 7(1):4.

35. Anderson PM: Glutamine- and N-acetylglutamate-dependent carbamoyl phosphate synthetase in elasmobranchs. Science 1980, 208(4441):291–293.

36. Anderson PM: Purification and properties of the glutamine- and N-acetyl-L-glutamate-dependent carbamoyl phosphate synthetase from liver of Squalus acanthias. J Biol Chem 1981, 256(23):12228–12238.

37. Sancho-Vaello E, Fernandez-Murga ML, Rubio V: Site-directed mutagenesis studies of acetylglutamate synthase delineate the site for the arginine inhibitor. FEBS Lett 2008, 582(7):1081–1086.

38. Brandman O, Ferrell JE, Jr., Li R, Meyer T: Interlinked fast and slow positive feedback loops drive reliable cell decisions. Science 2005, 310(5747):496–498.

39. Senkevitch E, Cabrera-Luque J, Morizono H, Caldovic L, Tuchman M: A novel biochemically salvageable animal model of hyperammonemia devoid of N-acetylglutamate synthase. Mol Genet Metab 2012, 106(2):160–168.

40. Heibel SK, Lopez GY, Panglao M, Sodha S, Marino-Ramirez L, Tuchman M, Caldovic L: Transcriptional regulation of N-acetylglutamate synthase. PLoS One 2012, 7(2):e29527.

41. Livak KJ, Schmittgen TD: Analysis of relative gene expression data using real-time quantitative PCR and the 2(-Delta Delta C(T)) Method. Methods 2001, 25(4):402–408.

42. Caldovic L, Lopez GY, Haskins N, Panglao M, Shi D, Morizono H, Tuchman M: Biochemical properties of recombinant human and mouse N-acetylglutamate synthase. Mol Genet Metab 2006, 87(3):226–232.

43. Qu Q, Morizono H, Shi D, Tuchman M, Caldovic L: A novel bifunctional N-acetylglutamate synthase-kinase from Xanthomonas campestris that is closely related to mammalian N-acetylglutamate synthase. BMC Biochem 2007, 8:4.

44. Morizono H, Caldovic L, Shi D, Tuchman M: Mammalian N-acetylglutamate synthase. Mol Genet Metab 2004, 81 Suppl 1:S4–11.

